# Home range is not constrained by numbers of hippocampal neurons

**DOI:** 10.1101/2025.10.08.681184

**Authors:** Mia C. Elbon, Greg Casalino, Suzana Herculano-Houzel

**Affiliations:** Dept. of Psychology, Vanderbilt Brain Institute, Vanderbilt University, Nashville TN

## Abstract

Numbers of hippocampal neurons vary by over three orders of magnitude across mammalian species. What evolutionary pressures shape this diversity? Given the role of the hippocampus in spatial mapping, the greater spatial navigation demands of larger home ranges may drive selection for more hippocampal neuron. Using data from 379 species, we crossed home range and population density data with cortical and hippocampal neuron counts predicted from clade-specific brain scaling laws to examine whether home range scales universally with estimated hippocampal neuron numbers across mammals. We confirm that home range scales universally with the inverse of population density across species and increases with body mass and metabolic rate. However, home range scaling with hippocampal or cortical neuron numbers differs by clade, such that bats, carnivorans and cetartiodactyls traverse home ranges over 1,000-fold larger than other mammals with equivalent hippocampal neuron numbers. These findings persist across data subsets controlling for study method, duration, and temporal scope. Numbers of hippocampal neurons are thus not limiting to spatial navigation in the wild, calling into question adaptationist explanations for the evolution of more hippocampal neurons based on a supposed need for increased spatial processing capacity. We propose that home range is determined primarily by population density, mediated by field metabolic rate and diet. The diversity in hippocampal neuron numbers across mammals, in turn, arises as a byproduct of clade-specific scaling of numbers of cortical neurons which we suggest is contingent on energetic opportunity, not on navigational or other cognitive demands.

**Significance Statement:** Why do some animals have so many more brain neurons than others? One explanation is evolution through selection for more neurons as needed for instance to navigate their environments to find food. However, this study finds that bats, carnivorans and whales traverse distances more than 1,000-fold larger than other mammals with similar numbers of neurons in the hippocampus, the brain structure required for spatial navigation. The authors question the traditional need-based explanation for brain diversity in favor of an opportunity-based account of the evolution of numbers of brain neurons.

## Introduction

There is enormous diversity in numbers of cortical neurons across mammalian species. What are the drivers of that diversity, that is, what explains why some species have very few and others have up to a thousand times as many cortical neurons? Ever since Darwin, need-based natural selection is as a rule invoked as the main driver of diversity in evolution. By that logic, and assuming that more neurons increase the information-processing capacity of brain structures, different parts of the brain with different functions should be found to have more neurons as species rely more heavily on those functions.

In what concerns the hippocampus, a cortical structure responsible for the associative learning underlying concept building and spatial navigation^1–6^ , those species with increasing needs for spatial navigation should be under selective pressure for increasing numbers of hippocampal neurons. Across mammalian species, the need for spatial navigation capacity should increase with home range, which is the area that an animal traverses over a set period of time in its normal activities of food gathering, mating, and caring for young^7^. Mammals create spatial maps using their hippocampus^1,8,9^It has been suggested that hippocampus size varies with relative selection pressures on cognitive mapping abilities and spatial memory^10–12^. Larger animals have increasingly larger home ranges^13–20^, which is considered to be a response to the larger metabolic needs of species with increasing body mass^7,13,14,18,19,20–24^. Increased metabolic needs would also explain the larger home ranges of carnivores compared to herbivores of similar body mass^13,19,24,25^. Accordingly, larger animals with larger home ranges should be under selective pressure for increased hippocampal volume. Indeed, animals who rely more on complex spatial behaviors (e.g., animals with larger home ranges) have larger hippocampi^10,26–30^. Similarly, increasing the spatial complexity of the environment has been shown to increase hippocampal volume^31–35^.

The underlying rationale behind the argument that animals need larger hippocampi to navigate larger home ranges is that larger hippocampi should consist of proportionately more neurons; absent direct data on numbers of hippocampal neurons, hippocampal volume has been expected to serve as a proxy for the number of neurons in the hippocampal network serving spatial navigation. So far, however, very little data exist on actual numbers of neurons in the hippocampus. Models shows that increasing numbers of neurons in the hippocampal network should afford a linear increase in the capacity of this network to represent the area of the environment that an individual traverses with a given spatial resolution^9,36–38^. If covering the increased surface areas of larger home ranges requires increased numbers of hippocampal neurons to the point that it imposes a selective pressure, then one should find a universal correlation of both hippocampal size and numbers of hippocampal neurons with home range across a wide variety of mammalian species.

We report in an accompanying paper that the total number of neurons in the mammalian hippocampus measured with the isotropic fractionator, a non-stereological method that provides estimates that are completely independent of structure mass, scales universally across species not with hippocampal or brain mass, but with the total number of neurons in the cerebral cortex^39^. Importantly, our finding of clade-specific scaling of numbers of hippocampal neurons with brain mass contradicts stereological estimates in a recent report that simply reflect brain size across a wide range of mammalian species^40^, which could be due to undersampling that makes stereological estimates inadvertently sensitive to structure size^41^. Our new finding that numbers of hippocampal neurons scale approximately with the square root of the total number of neurons in the cerebral cortex, both of which can be estimated from brain mass in a clade-specific manner, puts us in position to finally examine directly whether home range scales universally with the estimated number of hippocampal neurons across species covering the entire spectrum of mammalian body sizes. Here we use the home range dataset compiled by Broekman et al.^42^ to which we add estimates of population density for those same species compiled by Santini et al.^43^; data on body mass and basal metabolic rate compiled in the AnAge database by de Magalhães and Costa^44^; and data on field metabolic rates from three different sources^45–47^.We then cross these data both with numbers of neurons in the cerebral cortex predicted from brain mass in each species according to the clade-specific relationships that have been found to apply separately to primates, glires, certartiodactyls, afrotherians, marsupials, chiropterans, and carnivorans^48^; and with the numbers of neurons in the hippocampus of each species predicted from their number of cerebral cortical neurons using the scaling relationships that apply to their clade^39^. The latter was used as an added check because the universal scaling of numbers of hippocampal neurons with the square root of total numbers of cerebral cortical neurons^39^ means that any scaling relationships that apply to total numbers of cerebral cortical neurons also apply to hippocampal neurons with the exponent divided by 2. We could then test whether the expected universal correlation between home range and numbers of hippocampal neurons applied across mammalian species.

## Results

Across 350 mammalian species in our dataset, we find that average home range scales as power functions of adult body mass (Fig. 1A) and basal metabolic rate (Fig. 1B) with exponents 1.06 ± 0.04 and 1.36 ± 0.09, respectively. The residual variation in log_10_ average home range after accounting for log_10_ body mass or log_10_ basal metabolic rate shows that bats, cetaceans and most carnivorans, especially the semi-aquatic species and some bears (asterisks in Fig. 1), consistently have much larger home ranges than predicted for their body mass (Fig. 1C) and basal metabolic rate (Fig. 1D; ANOVA p<0.0001). However, the apparent effect of clade is due to diet type (Fig. 1E, F, both ANOVA p<0.0001), as carnivorous species have consistently larger home ranges than herbivorous and omnivorous species. Within each diet type, average home range scales similarly across clades (all ANOVA p>0.01, with the exclusion of bats, which are always outliers as shown in Fig .1A). Importantly, across herbivorous and omnivorous species (not including bats), average home range scales across clades with an exponent of 1.006±0.044 (r^2^=0.682, p<0.0001), identical to unity; in contrast, across carnivorous species (not including bats), average home range scales across species of all clades with a larger scaling exponent of 1.308±0.056 (r^2^=0.870, p<0.0001).

**Figure 1.**
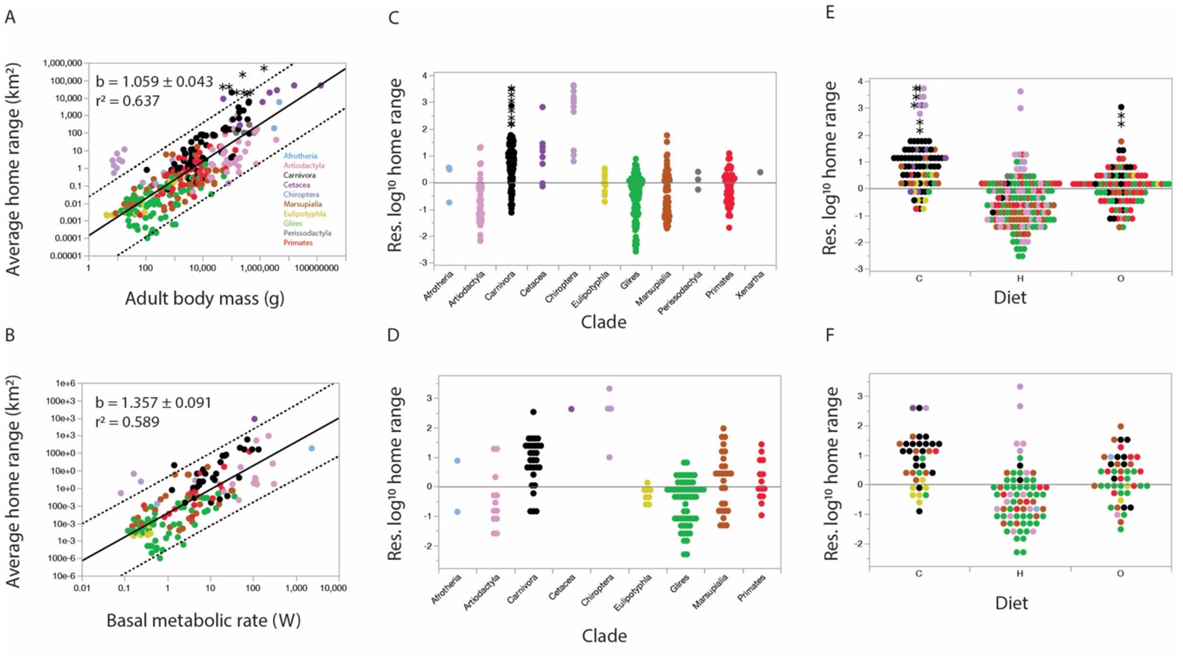
Average home range scales universally with body mass and basal metabolic rate across a wide range of mammalian species, with the exception of bats, semi-aquatic carnivorans and some bears. All plotted functions include all species and clades in the dataset. Allometric exponents and r^2^ values are shown in the graphs. **A**, average home range scales with adult body mass (p < 0.0001, *n =* 350) in a manner that leaves bats, semi-aquatic carnivorans and some bear species (asterisks) outside the 95% prediction interval (dotted lines). **B**, average home range scales with basal metabolic rate (p < 0.0001, *n =* 157) in an apparently universal fashion, although bats remain outliers; notice that data for aquatic carnivores, several bats and some bears are not available in the dataset. **C**,**E**, residual log10 home range for each species after accounting for log10 adult body mass, grouped by clade (**C**) or by diet type (**E**). **D**,**F**, residual log10 home range for each species after accounting for log10 basal metabolic rate, grouped by clade (**D**) or by diet type (**F**). Diet types are carnivory (C), herbivory (H) or omnivory (O).

In contrast, we find that the scaling relationships between home range and numbers of neurons in the cerebral cortex (Fig. 2A) and hippocampus (Fig. 2B) do not apply universally across all mammals or clades. Using the functions fitted across all species, primates consistently have smaller home ranges and cetaceans and chiropterans as well as semi-aquatic carnivorans consistently have larger home ranges than expected for their numbers of neurons, as the residuals show (Fig. 2C, D; ANOVA for all clades, p<0.0001; ANOVA for carnivorans, chiropterans, and primates, p<0.0001). Importantly, the distinct scaling of home range with numbers of cortical and hippocampal neurons in primates is not due to diet type (Fig. 2E, F), because primate home ranges remain significantly different (smaller than expected) after accounting for diet type (ANOVA for carnivorans, glires and primates, p<0.0001 for each diet type). Semi-aquatic carnivores (all of which are carnivoran species) and bats again are the species with the largest residual home range, now joined by several cetacean species (Fig. 2A, B). The systematic disparities between bats, carnivorans, cetaceans and primates suggest that the scaling of home range with numbers of cortical and hippocampal neurons is clade-specific.

**Figure 2:**
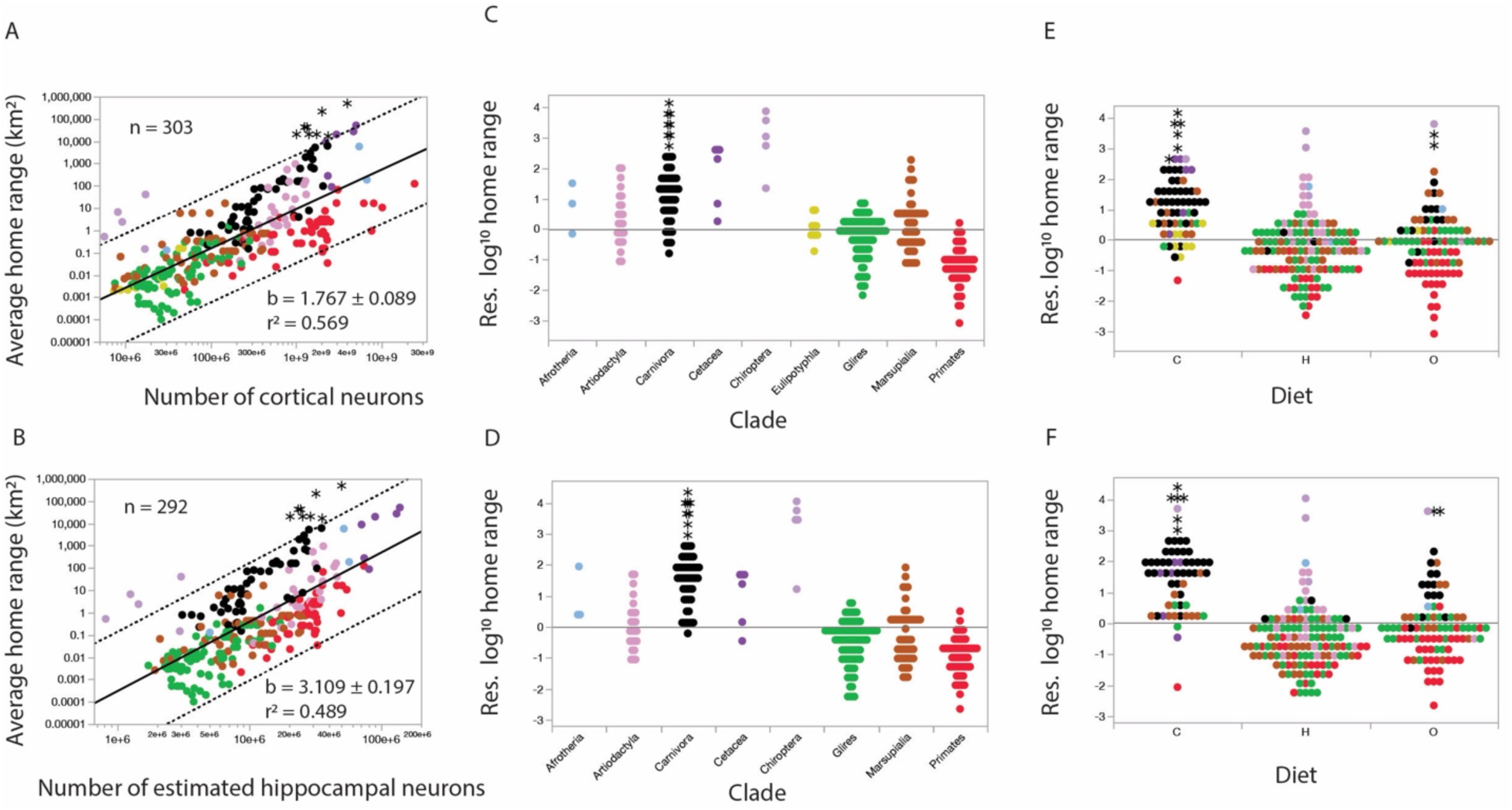
Home range does not scale universally with numbers of cortical or hippocampal neurons across mammalian species. All plotted functions include all species and clades in the dataset. Allometric exponents and r^2^ values are shown in the graphs. A, B, average home range scales with estimated total numbers of cerebral cortical neurons (A; p < 0.0001, *n =* 303) or of hippocampal neurons (B; p < 0.0001, *n =* 292) in a manner that leaves semi-aquatic carnivorans and some bear species (asterisks) as well as bats outside the 95% prediction interval (dotted lines), and additionally leaves all primate species below the fitted line, although mostly still inside the prediction interval. C, residual log_10_ home range for each species after accounting for log_10_ number of cerebral cortical neurons, grouped by clade. D, residual log_10_ home range for each species after accounting for log_10_ number of hippocampal, grouped by clade. Note that primate species consistently have smaller home ranges and carnivorans, especially the semi-aquatic species, consistently have larger home ranges than predicted for their numbers of neurons.

The clade-specificity of the scaling of home range with numbers of cortical and hippocampal neurons is evidenced in Figure 3. Glires, eulipotyphlans, marsupials, afrotherians and primates share scaling exponents below 3.0 or 6.0 (for numbers of cortical or hippocampal neurons, respectively), whereas carnivorans and artiodactyls (including cetaceans) share exponents above those values (exponents indicated in Fig. 3A, B; the scaling function for chiropterans is not significant, with p=0.7757). Interestingly, while bats are always outliers with much larger home ranges for their numbers of neurons, as with body mass (Fig. 1), the data for the remaining clades can be grouped in two different ways that make either primates or large carnivorans and cetartiodactyls into outliers. When carnivorans and cetartiodactyls (in addition to bats) are excluded from the analysis, yielding joint power functions with an r^2^ of 0.641, primates are well aligned with glires, marsupials and eulipotyphlans in the scaling of home range with numbers of neurons (Fig. 3A, B). In this case, all large carnivorans (not only the semi-aquatic species) and cetartiodactyls would be described as having far larger home ranges than predicted for their numbers of cortical and hippocampal neurons. Alternatively, if only primates (and bats) are excluded while carnivorans and cetartiodactyls are included in the analysis together with all other clades (Fig. 3C, D), we find that average home range scales with numbers of all cortical or hippocampal neurons as power functions of a higher r^2^ of 0.788 and 0.655 that however still exclude the aquatic carnivorans and some bears in the dataset, as well as bats. Using this latter analysis, which provides a better description of the variation in the dataset, primates and bats are the outlier clades, and in different directions: primates consistently occupy much smaller home ranges, and most bats consistently occupy much larger home ranges, than other mammals of similar numbers of cortical or hippocampal neurons. Either way, two facts remain: that home range and numbers of cortical and hippocampal neurons do not vary in a universally correlated manner across mammalian species; and that semi-aquatic carnivorans, many bats and some bears occupy much larger home ranges than can be predicted by their body size or numbers of neurons.

**Figure 3.**
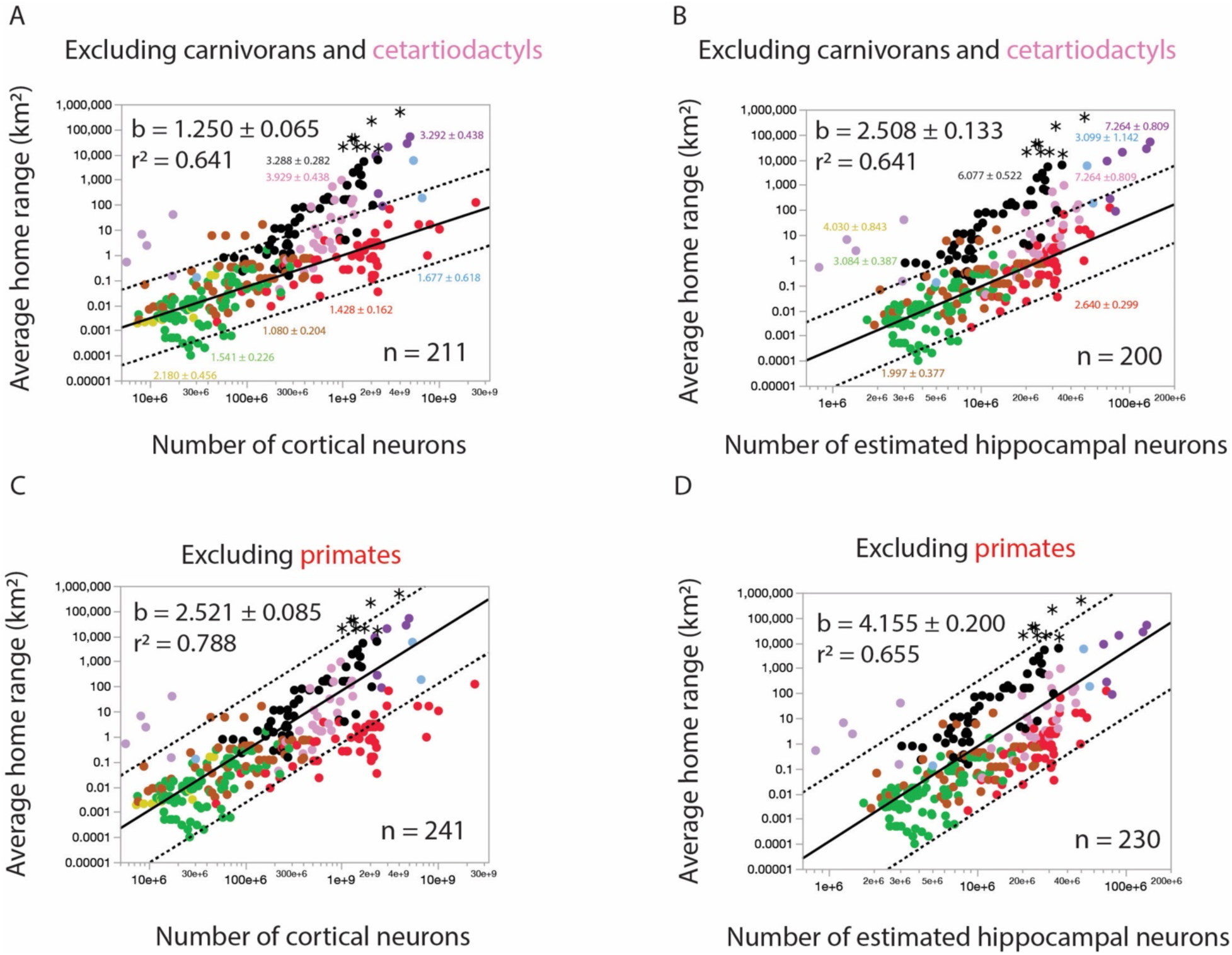
Home range scales with numbers of cortical or hippocampal neurons in a clade-specific manner. Scaling exponents ± standard error that apply to each clade (not plotted) are indicated in the graphs. Plotted functions include all species and clades in the dataset (excluding chiropterans) minus carnivorans and cetartiodactyls (A and B) or minus primates (C and D). Allometric exponents and r^2^ values for the plotted functions are shown in black in the top left of each graph. **A and B:** When carnivorans and cetartiodactyls are excluded from analysis, average home range scales with number of cortical neurons and number of hippocampal neurons (p < 0.0001, *n =* 216) in a manner that includes primates and leaves most carnivorans, most cetaceans and some artiodactyls outside the 95% prediction interval (dotted lines). **C and D**: When primates are excluded from analysis, average home range scales as power functions of numbers of cortical and hippocampal neurons (p < 0.0001, *n =* 216) with a higher r^2^ and a 95% prediction interval that either excludes primates (C) or leaves them systematically below the prediction line (D) and still excludes the semi-aquatic carnivorans and some bears in the dataset. Notice that most bats in the dataset are always outliers.

### Effect of tracking methods, sample size, and study duration

The average home range estimates used so far in the analysis combine various tracking methods, sample size, and study duration, as has been standard practice in the field^19,42,50,51^. The lack of consistence in the time span (daily, annual, other) and level (individual, group, population) that apply to the values of home range analyzed raises the concern that the finding that primates are outliers in the scaling of home range with numbers of neurons is a measurement artifact. To address this possibility, we determined whether primates are still outliers when the scaling between numbers of cortical or hippocampal neurons and home range is calculated exclusively for subsets of data restricted to the same combination of time span (home range covered in a day or in a year) and level (home range of single individuals or of a population). Figure 4 shows that even in these much more restricted datasets (which lack data for bats, semi-aquatic carnivorans and cetaaceans), and despite the inclusion of primates in the fitted data, species in this clade still consistently have smaller home ranges than predicted for their numbers of cortical or hippocampal neurons, well below the 95% prediction interval.

**Figure 4.**
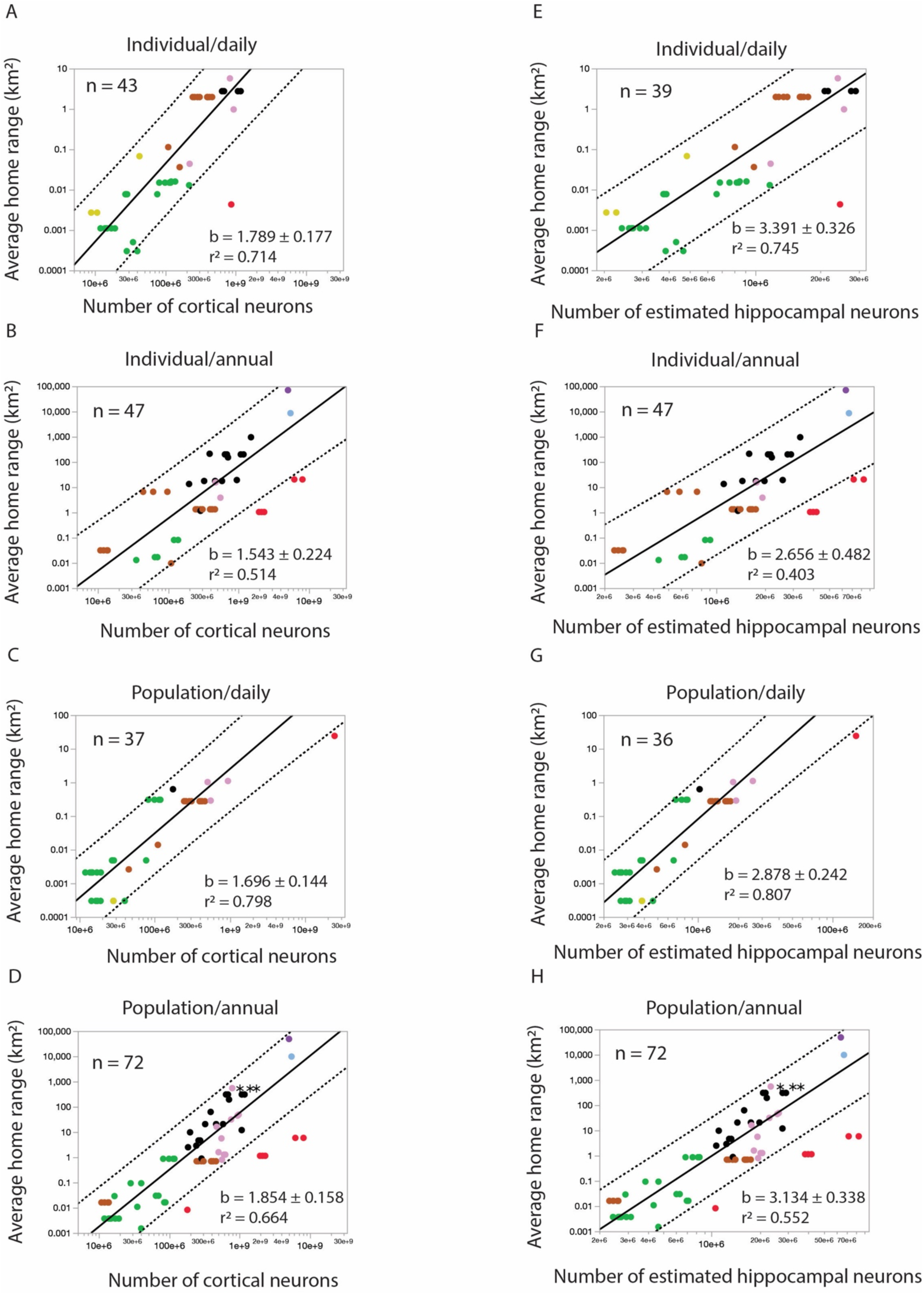
Primates have smaller home ranges than predicted for their numbers of cortical and hippocampal neurons even in data sets restricted to similar time span and population level. Average home range is plotted as a function of numbers of cortical neurons (A-D) or estimated numbers of hippocampal neurons (E-H) using data restricted to daily home range measured in individuals (A, E); annual home range measured in individuals (B, F); daily home range measured in a population (C, G); or annual home range measured in a population (D, H). The plotted functions apply to all species in each data set, including primates, which still fall below the 95% prediction interval (dotted lines) in every case. Allometric exponents (all p<0.0001), r^2^ values and n are shown in the graphs.

### Population density is a universal predictor of home range, and also does not scale universally with numbers of cortical neurons

We find that average home range scales with the near inverse of population density, with an exponent indistinguishable from -1 (Fig. 5A), in a manner that applies equally across carnivores, herbivores and omnivores (Fig. 5B, ANOVA p=0.3789) and, importantly, also includes bats (Fig. 5C). Our dataset thus reproduces the previous observations that home range is a negative inverse function of population density^15,18,52–54^. While an ANOVA across all clades (Fig. 5C) indicates a significant effect of clade (p<0.0001), residual home ranges after accounting for population density do not differ across the three clades that share large numbers of hippocampal neurons but differ markedly in average home range (ANOVA for carnivorans, cetartiodactyls, and primates, p=0.1429), nor across bats and rodents that share small numbers of hippocampal neurons but also differ strikingly in home range (ANOVA, p=0.3347). Interestingly, while the scaling of home range with the inverse of population density includes bats, primates and cetaceans, most semi-aquatic carnivore species remain outliers, with larger home ranges than predicted for their population density (Fig. 5A).

**Figure 5.**
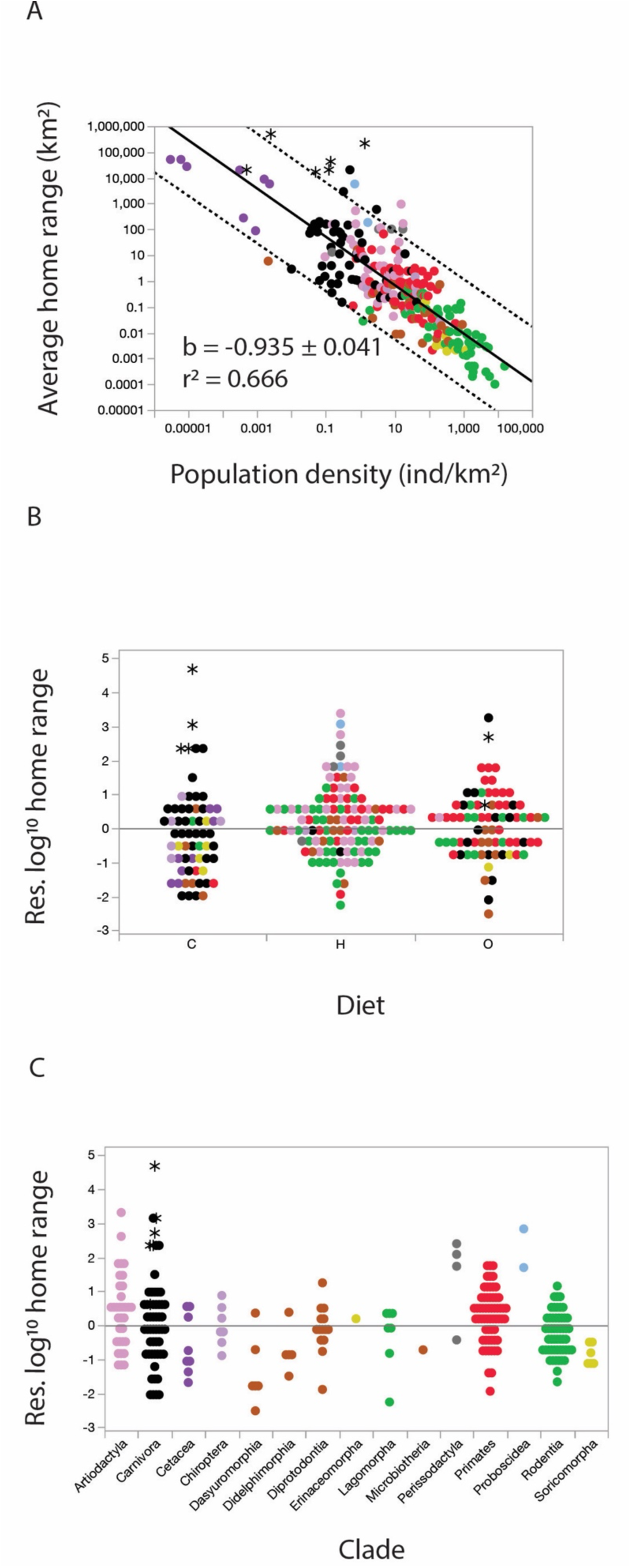
Population density is a better universal predictor of home range than any other variable examined. A, home range scales as a power function of population density (A, p < 0.0001, *n =* 262). B, residual log_10_ home range by population density for each species grouped by diet. C residual log_10_ home range by population density for each species grouped by clade.

In line with the lack of a universal correlation between home range and numbers of hippocampal neurons, we find that the power functions that relate population density to numbers of cortical (Fig. 6A) or hippocampal neurons (Fig. 6B) are not universal across all mammals in the dataset (both ANOVA p<0.0001), and also do not apply equally across species of different diet types (Fig. 6C,D, ANOVA p<0.0001). Across clades primates and glires have systematically higher and carnivorans, cetaceans and the one bat species with estimates available have systematically lower population densities than predicted for their numbers of neurons (Fig. 6E,F), even when analysis is restricted to herbivorous and omnivorous diet types (ANOVA, p<0.0001), which argues for clade-specific relationships between population density and numbers of hippocampal and cortical neurons.

**Figure 6.**
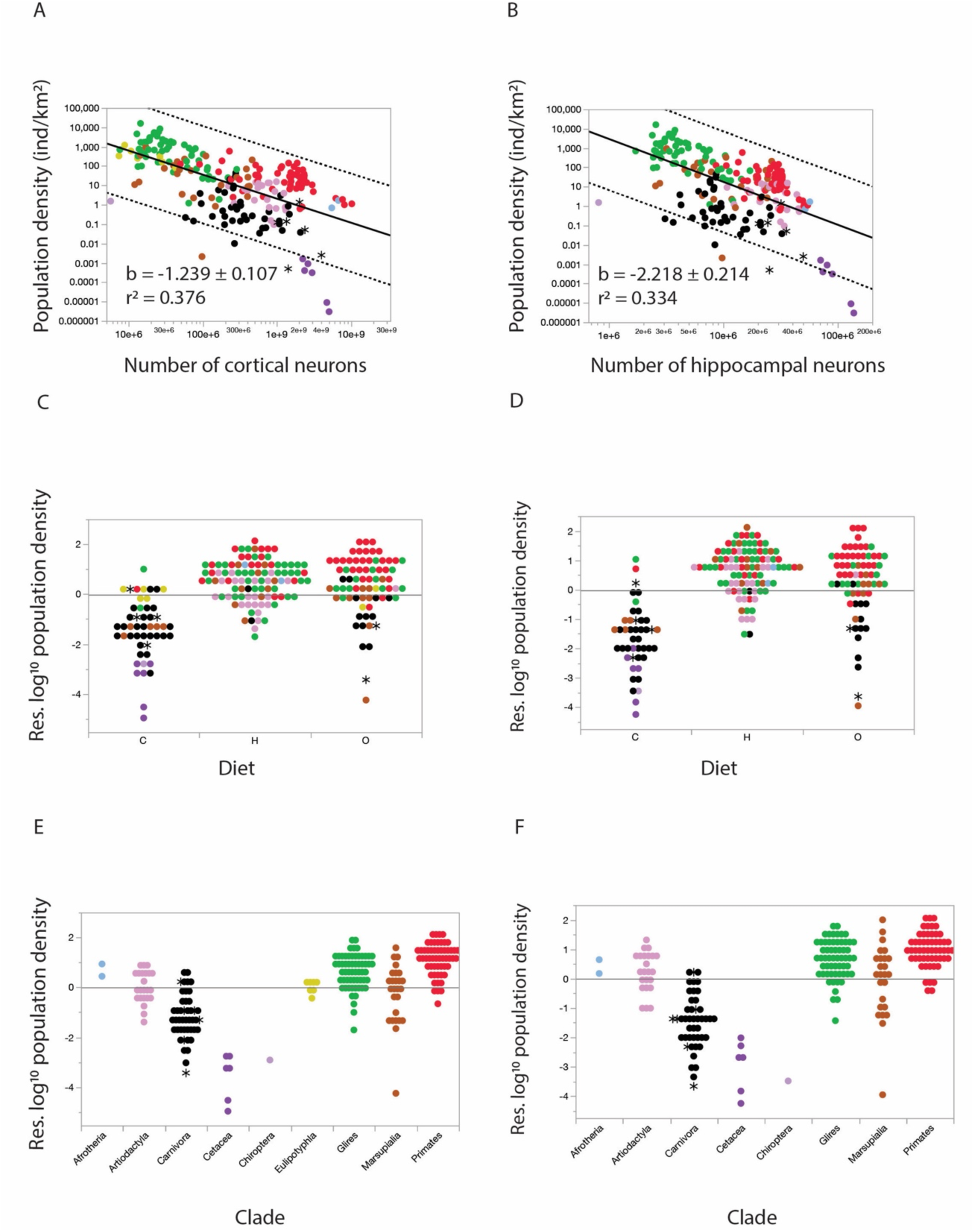
Population density is not a universal predictor of number of hippocampal neurons. Population density scales as a power function of number of cerebral cortical (A; p<0.0001. n=XXX) or hippocampal neurons (B; p < 0.0001, *n =* 216) across species in a manner that includes primates but still leaves cetaceans and one semi-aquatic carnivoran (asterisks) below the 95% prediction interval (dotted lines). C, D, residual log_10_ population density after accounting for number of cerebral cortical (C) or hippocampal (D) neurons for each species, grouped by diet type. E, F, residual log_10_ population density after accounting for numbers of cortical (E) or hippocampal (F) neurons for each species, grouped by clade. Note that there is a single species of chiropterans with data available for both population density and estimated numbers of neurons.

Finally, we sought to determine the source of the larger home ranges and lower population densities of carnivorans and carnivorous species as a whole compared to primates of similar numbers of hippocampal neurons by examining how these three variables scale with the field metabolic rate that indicates how much energy different species must be able to procure per day in order to survive. Figure 7 shows that field metabolic rate is a good but not universal predictor of home range (Fig. 7A), population density (Fig. 7B) and number of hippocampal neurons (Fig. 7C). Carnivorous species (Fig. 7D) and also bats and carnivoran species (Fig. 7G, the vast majority of which also have carnivorous diets) consistently have larger home ranges, whereas primates and glires have consistently smaller home ranges (Fig. 7G), than predicted for their field metabolic rate. Carnivorous species, which include carnivorans and cetaceans and the one available bat species, also have lower than predicted population densities, while primates have higher than predicted population densities, for their field metabolic rates (Figs. 7E, H). Diet, on the other hand, does not differentiate the relationship between numbers of hippocampal neurons and field metabolic rate (Fig. 7I; ANOVA, p=0.0343 versus p<0.001 for all other comparisons); the distinguishing factor here is clade, as primates have systematically more hippocampal neurons than predicted for their field metabolic rates (Fig. 7F), in line with their systematically larger numbers of cortical neurons compared to non-primate mammals of similar body mass^50^, and the two bats with data available have fewer hippocampal neurons than predicted for their field metabolic rates. We thus find that while the home range occupied by mammalian species is a universal function of their population density, with both variables affected coordinately by diet, the number of neurons in the hippocampus of mammalian species is not universally related to the home range, population density, or field metabolic needs of a species.

**Figure 7.**
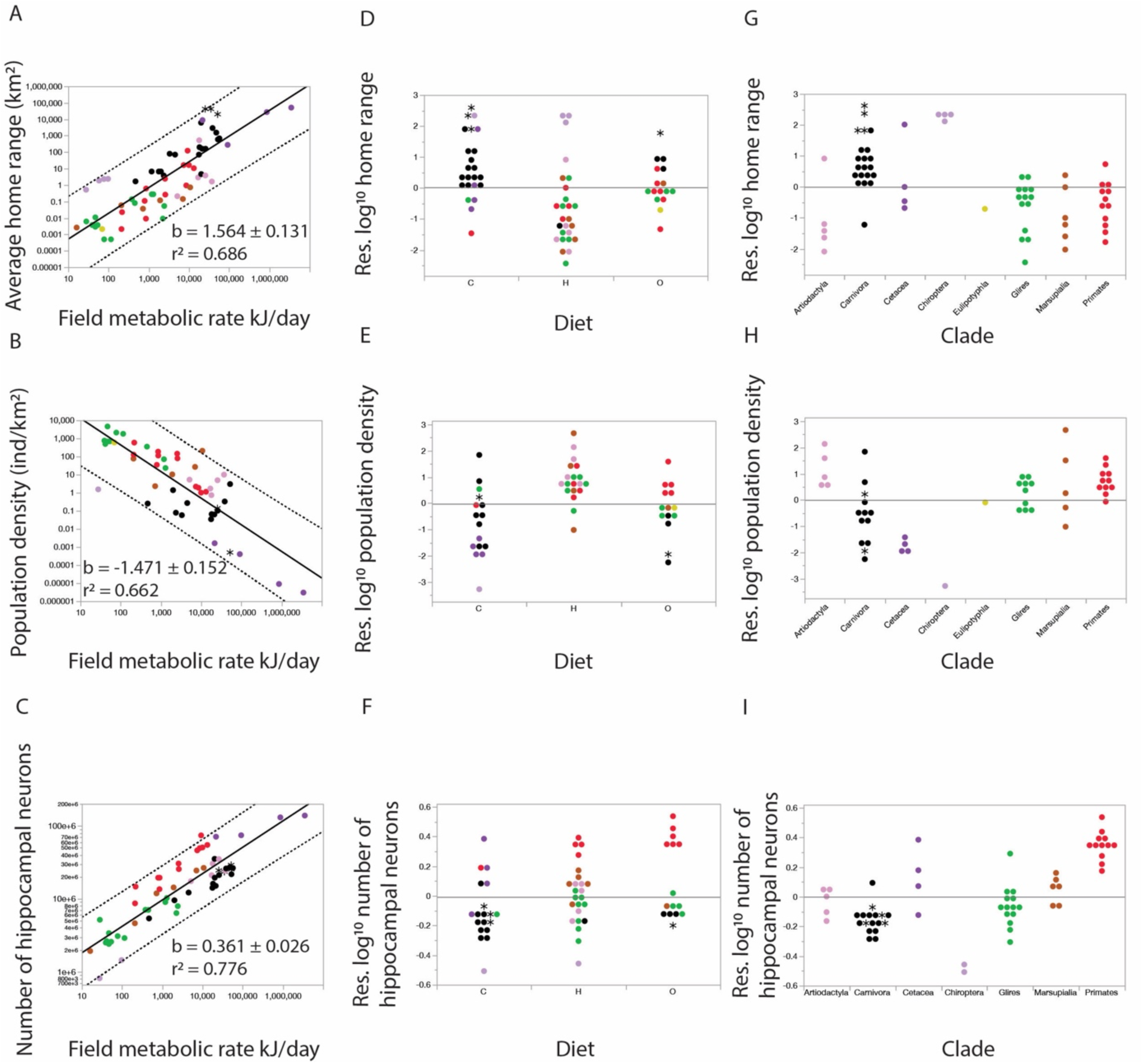
Field metabolic rate mediates population density. Field metabolic rate scales as a power function of average home range (A), population density (B), and estimated number of hippocampal neurons (C) across mammalian species. All plotted functions include all species and clades in the dataset. Allometric exponents and r^2^ values are shown in the graphs. A-C, average home range scales with field metabolic rate (A; p < 0.0001, *n =* 67), population density (B; p < 0.0001, *n =* 50), and number of hippocampal neurons (C; p < 0.0001, *n =* 60) in a manner that includes semi-aquatic carnivorans and some bear species (asterisks) and primates inside the 95% prediction interval (dotted lines). D-I, residual log_10_ home range, population density, and number of hippocampal neurons after accounting for field metabolic rate for each species, grouped by clade (D-F) or by diet (G-I). Note that while primates are included in the 95% prediction interval, they deviate systematically from the predicted values.

## Discussion

Obtaining sufficient energy to make it repeatedly through the days, together with maintaining physical integrity, is the one non-negotiable condition for animal survival. Our findings are consistent with a scenario in which, given an environmental reality of limited food availability per surface area, the higher rates of energy use per day of larger animals cause them to self-organize, through food-seeking exploitative behaviors, into predictably lower population densities, which in turn necessitate exploratory behaviors and dispersal into inversely larger home ranges^15^. Importantly, our finding that home range scales linearly with body mass across herbivorous and omnivorous species but with a larger exponent of 1.3 across carnivorous species indicates that the inconsistent scaling exponents reported in the literature were due to the inclusion of varying numbers of carnivorous species (^14,17–19,22,25,49^). We propose that diet type affects feeding efficacy and efficiency and thus modulates the relationship between field metabolic rate and population density (Fig. 7), which, given the necessarily inverse relationship between population density and home range^20^, explains the dependence on diet of the scaling of home range with body mass^19^. Specifically, carnivory exposes animals to the relative scarcity of prey compared to vegetable foodstuffs per surface area in the environment^18,19,54^. As a result, for a given body size and field metabolic rate, carnivorous animals exist in lower population densities and explore larger home ranges than omnivorous animals, and herbivorous species exist in larger population densities and explores smaller home ranges, all the while home range and population density continue to scale universally across all diet types.

While we find that microchiropterans have larger home ranges than expected for their body mass, micro- and megachiropteran bats alike have the home ranges predicted for their population densities, which is explained by unusually low population densities for the body mass found in microchiropterans. Low population densities could be expected for insect-eating bats due to food scarcity combined with their very high specific metabolic rates, even if two of these bat species with very large home ranges in the dataset also consume fruits (*Carollia perspicillata* and *Glossophaga soricina*). Similarly, we propose that the relative scarcity of prey combined with the increased metabolic cost of living in cold water and the three-dimensional nature of hunting in an aquatic environment explain the finding that semi-aquatic carnivorans and cetaceans are consistently outliers in our dataset, exploring home ranges that are larger than expected not only for their body mass and field metabolic rate but also population density. The resting metabolic costs of aquatic mammals are about twice as high as those of terrestrial mammals, with a scaling relationship for resting metabolic rate with body mass of 26.9 M_bd_^0.69^ in aquatic mammals^56^ in contrast to McNab’s allometric prediction for the basal metabolic rate of similarly sized terrestrial mammals^57^ of 10.55 M_bd_^0.724^ (after converting for M_bd_ in kg), probably due to heat lost to cold water. The effect of prey scarcity to drive decreased population density and larger home range is expected to be especially severe in both aerial and aquatic environments, where there is an added dimension to prey movement. Interestingly, given the difference between stroke vs. stride energetic costs, fully submerged mammals are able to avoid hydrodynamic drag and exhibit lower mean stroke costs compared to surface swimming carnivores^58^. The higher efficiency of fully submerged carnivores, such as cetaceans, might explain how come their home ranges match the predicted values for their very low population densities whereas the aquatic carnivorans, as inefficient surface swimmers, explore much larger home ranges than predicted for their population densities (Fig. 5A).

To the heart of our central question, we find that the enormous range of coordinated variation in home range and population density occurs without the expected universal correlation with numbers of hippocampal neurons typically presumed to be necessary to support spatial exploration^10,26–35^. To the contrary, we show that carnivoran species are capable of exploring home ranges of around 1,000 km^2^ on land and as much as 100,000 km^2^ over water with the same estimated ca. 30 million hippocampal neurons found in primates who remain in an area of just about 1 km^2^; likewise, bats explore areas between 1 and 10 km^2^ with fewer than the 2-3 million hippocampal neurons found in small rodents and marsupials who forage in under 0.01 km^2^ (Fig. 2B). Conversely, bats, carnivorans and primates that explore home ranges of about 10 km^2^ do so with numbers of hippocampal neurons that are 10 times as large in primates as in carnivorans and ca. 30 times as large as in bats. If bats, carnivorans and primates alike obtain the energy that they need by occupying the home range that is predicted for their population density (Fig. 5A), the number of hippocampal neurons available to represent the surface area covered by a species is clearly not limiting to the point that it would be under selective pressure to increase together with home range as animals became larger in evolution. Known mechanisms of multiplexing hippocampal representations of environments on different scales^38^ and dynamically scaling place field sizes to context^59^ would suffice to account for how a given number of hippocampal neurons suffices to allow exploration of home ranges varying by a factor of over 1,000-fold that we uncover here.

The finding that bats cover inordinately large home ranges for their diminutive body mass using the smallest numbers of hippocampal neurons in our dataset should suffice to dismiss the notion that a need to navigate large environments imposes a selective pressure for more neurons, or even larger brains for that matter. Those determined to find adaptationist explanations to the discrepancy in scaling of home range with numbers of hippocampal neurons between primates and other mammals might turn to Fig. 3D and argue that non-primate species with larger home ranges do have the correspondingly larger numbers of hippocampal neurons predicted by a supposed need for more hippocampal spatial representational power. In that case, however, a new argument would have to be invoked to explain how come primates evolved much larger numbers of hippocampal neurons than “necessary” to represent their comparatively small environments – for instance, that primates benefit from the increased complexity in representing their environment afforded by more neurons (see below). In our view, a much more parsimonious explanation lies in acknowledging that the scaling relationship between numbers of neurons and body size is not universal^55,60^. If there is any correlation between home range and numbers of hippocampal (and cerebral cortical) neurons, we argue that it is simply because larger body mass, which necessitates universally larger home ranges as predicted by their population densities (according to the combination of field metabolic rate and diet), come with more cortical neurons^61^ – but following scaling rules that differ between primate and non-primate clades^55^. In this case, the clade-specific correlation between home range and numbers of hippocampal neurons is a trivial mathematical contingency, not a biological necessity.

This is not to say that larger numbers of hippocampal neurons do not bring a cognitive advantage. According to theory, the more the neurons in the hippocampal network, the bigger should be the capacity of the network to represent an area with a certain spatial resolution^38^, which can be presumed to extend to cognitive representations. Given the combination of the predictably small home ranges of primate species and their large numbers of hippocampal neurons (which are also predictable for their cerebral cortices^39^), we propose that primates do benefit from both increased complexity in cognitive representations and increased spatial resolution in their hippocampal maps of the environment, as the radius of their place cells is expected to be reduced^38^. This latter prediction can be tested with systematic comparisons of place cell radius across primate and non-primate species of either similar home ranges or numbers of hippocampal neurons. Evidently, the comparatively lower spatial resolution of non-primates and presumably simpler cognitive representations are perfectly sufficient for their successful navigation in exploring and exploiting whatever size home range is required for their survival. Thus, the increased spatial resolution we predict for primates cannot be considered a necessity, and therefore also not the result of selective pressures – unless one resorts to a convoluted argument that only primates were subject to such pressures for increased spatial resolution.

Rather, we propose that it is more parsimonious that in exploring their environment, mammals go as far as they have to go in order to meet their metabolic demands (given how their diet impacts population density), but their numbers of hippocampal neurons are just not limiting to how far they can go. The diversity of numbers of hippocampal neurons, which cannot be said to have been driven by selective pressure for increased spatial navigation capacity, is tied to the square root of the variation in total numbers of cortical neurons^39^. Thus, what needs to be explained is what leads to the enormous variation in total numbers of cortical neurons in mammalian evolution. We favor the simpler alternative explanation that numbers of cortical (and so also hippocampal) neurons increase in evolution as consistently increased energetic opportunities present themselves to a population across generations^62^, with body size increasing in tandem as still affordable^61^.

## Materials and Methods

The home range and population densities of 342 mammalian species were compiled using two databases: “HomeRange: A global database of mammalian home ranges"^42^ and “TetraDENSITY 2.0 – A database of population density estimates in Tetrapods”^43^. Home range data collection and statistical techniques for each species differed across studies, with often differing temporal measurements and differing observation techniques employed for any given species. To combine home range and population density data into a single database, it was necessary to calculate average home range values for each species. The combined database was then integrated with data on body mass, basal metabolic rate and diet type compiled by de Magalhães and Costa^44^ in the AnAge database and with data on numbers of cortical neurons predicted from brain mass^48^. From the numbers of cortical neurons, the number of hippocampal neurons was estimated for each species according to clade-specific formulas: e^7.673^ N_cx_^0.815^ for cetartiodactyls, e^3.181^ N_cx_^0.658^ for carnivorans, e^-4.792^ N_cx_^1.183^ for chiropterans, e^4.322^ N_cx_^0.640^ for marsupials, e^9.841^ N_cx_^0.346^ for primates, e^3.384^ N_cx_^0.693^ for glires, and e^7.767^ N_cx_^0.450^ for afrotherians. Note that these clade-specific calculations provide an alternate set of estimates for N_hp_, which can also be more simply obtained as roughly N_cx_^0.5^ [39]. Finally, field metabolic rate data were added to the combined database using values gathered from Nagy^45^, de Castro^46^ and Pontzer^47^.

From these data, average home range (measured in km^2^) for each species was plotted in log-transformed bivariate-fit graphs comparing home range against adult body mass (measured in grams), basal metabolic rate (in W), field metabolic rate (in kJ/day), number of cortical neurons (N_cx_), estimated number of hippocampal neurons (N_hp_), and population density (individuals/km^2^) in JMP Pro 18. Mammalian data points were separated by clade for comparison, as indicated in the graphs. We purposefully do not use any phylogenetic correction methods because clade effects are addressed directly in our analysis.

## Acknowledgments

Thanks to Vanderbilt University for supporting Mia C. Elbon. This work was not supported by any external sources of funding.

## Notes

### Competing Interest Statement

The authors have declared no competing interest.

### Summary of Updates

Added additional data for Chiroptera; addressed recent workin the literature

